# Persistence of Cajal-Retzius cells in the postnatal hippocampus is required for development of dendritic spines of CA1 pyramidal cells

**DOI:** 10.1101/2022.05.09.491146

**Authors:** Ingvild Lynneberg Glærum, Keagan Dunville, Nicola Pietro Montaldo, Hinako Kirikae, Maximiliano Jose Nigro, Pål Sætrom, Barbara van Loon, Giulia Quattrocolo

## Abstract

Cajal-Retzius (CR) cells are a transient type of neuron that populate the postnatal hippocampus. The role of transient cell types and circuits have been vastly addressed in neocortical regions, but poorly studied in the hippocampus. To understand how CR cells’ persistence influences the maturation of hippocampal circuits, we specifically ablated CR cells from the postnatal hippocampus. Our results highlighted layer-specific effects on dendritic spines and synaptic-related genes and revealed a critical role of CR cells in the establishment of the hippocampal network.

## Introduction

The hippocampal formation is a brain structure primarily associated with memory formation, learning and spatial representation (Buzsáki & Moser, 2013). The microcircuits responsible for these functions are refined around birth and during the first few weeks of postnatal development (Alberini & Travaglia, 2017; Donato et al., 2017; Dumas, 2005; Travaglia et al., 2016). At this time, hippocampal neurons undergo changes in morphology, physiological properties and connectivity patterns. These changes are guided by unique genetic programs (Yuste et al., 2020).

Cajal-Retzius (CR) cells are reelin-expressing glutamatergic neurons, whose function has mainly been correlated to the orchestration of cortical lamination in prenatal development (D’Arcangelo et al., 1995; Gil et al., 2014; Ramón y Cajal et al., 2011). Although most neocortical CR cells disappear soon after birth, 20% of hippocampal CR cells persist for several months (Anstötz et al., 2016; Ma et al., 2014; Supèr et al., 1998). In the hippocampus, CR cells are localized in the outer molecular layer of the dentate gyrus and the stratum-lacunosum moleculare (SLM) of the CA-regions (Anstötz et al., 2016; Del Río et al., 1997). The axons of CR cells have extensive axon collaterals that can occasionally be long ranging, terminating as far as the entorhinal cortex (Ceranik et al., 1999; Deng et al., 2007). The long-range projections of CR cells have been proposed to serve as a scaffold for entorhinal fiber innervation of the hippocampus, with CR cells serving as transient targets of entorhinal afferents (Anstötz et al., 2016; Deng et al., 2007; Supèr et al., 1998).

While CR cells seem to have a clear role during prenatal development, little is known about their function in postnatal hippocampal development. Electrophysiological studies have shown that CR cells are actively integrated in the hippocampal microcircuit, receiving GABAergic inputs, and generating glutamatergic outputs to both pyramidal neurons and inhibitory interneurons (Anstötz et al., 2018; Quattrocolo & Maccaferri, 2013; Quattrocolo & Maccaferri, 2014), though their impact on the maturation of the hippocampal circuits remains poorly understood.

To determine the influence of hippocampal CR cells on postnatal circuit development, we established a model by which we can successfully ablate CR cells from the postnatal hippocampus. By combining spine analysis, mRNA sequencing, and electrophysiology we identified significant and layer-specific changes in dendritic spines and in the expression of synaptic-related genes. Our findings indicate that the persistence of CR cells in the postnatal hippocampus is critical for proper maturation of the hippocampal circuit.

## Results

### Targeting and ablation of hippocampal Cajal-Retzius cells

To determine the role of CR cells in the developing hippocampus we took advantage of the Pde1c-Cre transgenic mouse line which conditionally expresses Cre in CR cells (Osheroff & Hatten, 2009). To confirm the validity of the transgenic line for the hippocampus, we crossed a Pde1c-Cre^+/-^ mouse with a *Ai9* TdTomato reporter mouse and performed immunohistochemical labeling for CR-cells specific markers, reelin and p73 (Meyer et al., 2004). Nearly all tomato^+^ cells expressed reelin and p73, with no significant difference between the cell ratio (Figure 1A-C). Thus, the Pde1c-Cre transgenic mouse line successfully labeled hippocampal CR cells.

**Figure 1.**
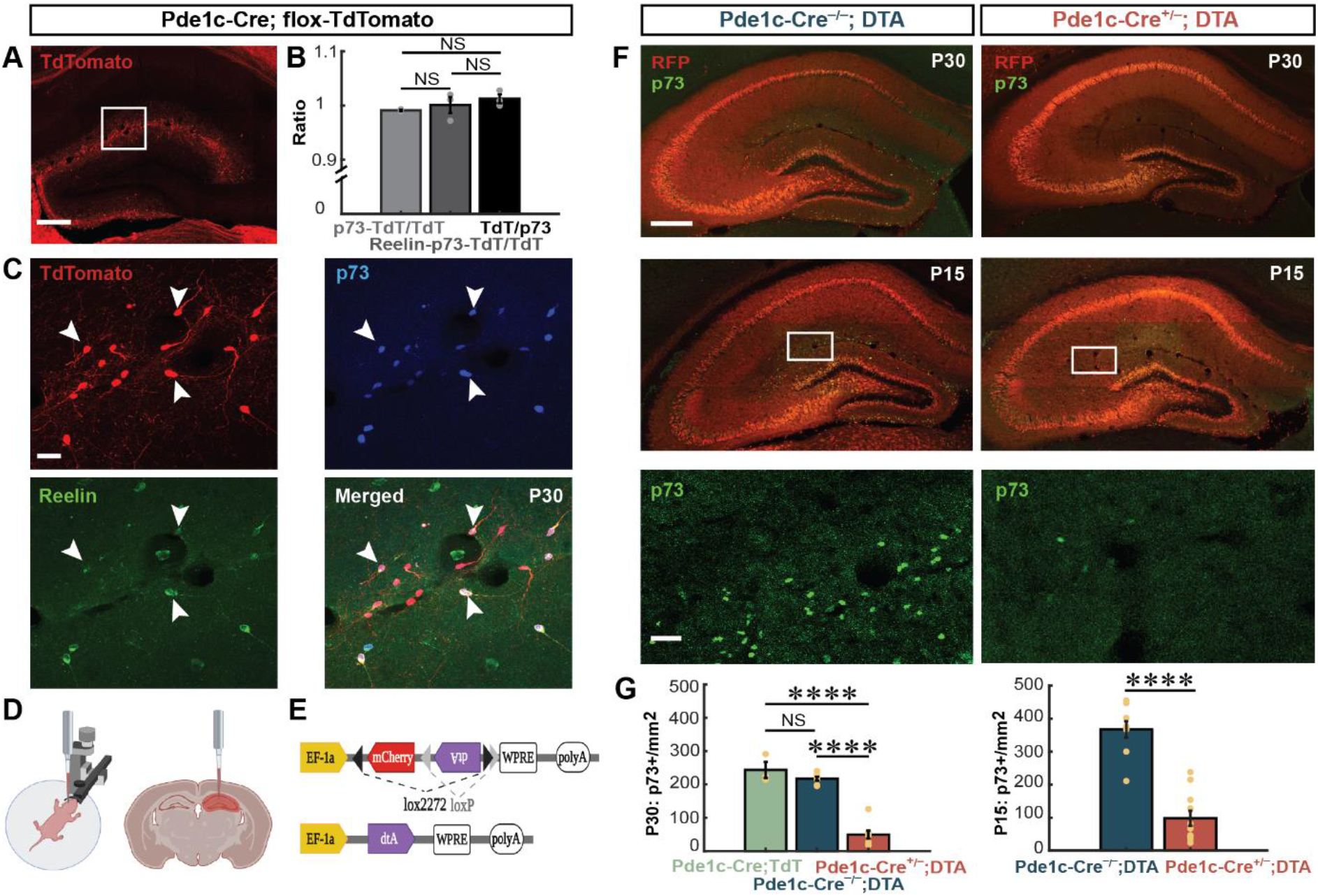
Targeting and ablation of hippocampal Cajal-Retzius cells. (A) Image of coronal section depicting tomato^+^ cells in the hippocampus of a Pde1c-Cre^+/-^;flox-TdTomato. (B) Bar graph illustrating ratio of counted immunolabelled p73+, TdTomato^+^, and Reelin+ cells (one-way ANOVA, n=3 animals). (C) Magnified insets (*white square)* from A depicting immunolabelled TdTomato^+^, p73^+^, and Reelin^+^ cells. (D) Schematic illustrating pup injection procedure. (E) Schematic of injected virus structure. (F) Examples from P30 (*top panels*) and P15 (*middle panels*) of Pde1c-Cre mice injected with AAV-DTA virus. Bottom panels represent high-magnification image of the inset from middle panels. (G) Comparative analysis of p73^+^ cell densities in Pde1c-Cre^+/-^;flox-TdTomato (n=3), Pde1c-Cre^-/-^;DTA (P30,n=7;P15,n=9) and Pde1c-Cre^+/-^;DTA samples (P30,n=11; P15,n=8) at P30 (*left*) and at P15 *(right)* using two-sample t-tests. Bar graphs show group mean ± SEM. Scale bar in (A) and (F) 200 µm, magnified insets in (C) 20 µm and (F) 50 µm. p-values: NS ≥ 0.05, **** ≤ 0.0001

To study the contribution of CR cells to the maturation of the postnatal hippocampus, we prematurely induced their death. At P0, we injected bilaterally in the dorsal hippocampus of Pde1c-Cre mouse pups a viral vector (pAAV-mCherry-flex-dta) (Wu et al., 2014), encoding mCherry (panneuronaly) and diphteria-toxin fragment A (DTA) (Cre-dependently) (Figure 1E). This construct drives the expression of mCherry in all the infected cells, allowing for an easy visualization of the injected site, while only Cre-expressing cells express DTA and undergo apoptosis. Immunostaining for p73 showed a 73% reduction in the number of hippocampal CR cells in Pde1c-Cre^+/-^;DTA at P15 and a 77% reduction at P30 (Figure 1F,G). CR cell density in control animals was similar to that observed in Pde1c-Cre^+/-^;flox-TdTomato mice, suggesting the injection procedure did not directly affect the survival of CR cells (Figure 1G). These results indicate that our targeted approach drastically reduces the postnatal density of CR cells and can thus provide insights on the consequences of their ablation on the development of the hippocampal circuit.

### Ablation of CR cells disrupts the formation of dendritic spines on CA1 pyramidal cells

We then sought to investigate the impact of CR cell ablation on CA1 pyramidal cells, the principal output neurons of the hippocampus and postsynaptic targets of CR cells (Quattrocolo & Maccaferri, 2014). Prenatal reduction of CR cells in the barrel cortex reduces spine density in apical dendrites of cortical pyramidal cells, whereas preventing the death of CR cells in the postnatal somatosensory cortex caused an increase in spines on both apical and basal dendrites (de Frutos et al., 2016; Riva et al., 2019). Therefore, we started by performing a morphological assessment of dendritic spines at juvenile stages in CR cells-ablated mice.

First, we quantified and classified spines present in terminal dendritic segments (Figure S1B) in the SLM (Figure 2A), Stratum Radiatum (SR) (Figure 2C) and Stratum Oriens (SO) (Figure S1C). We observed no difference in the total number of spines in the SLM, but a small significant reduction of dendritic spines in SR and SO after CR cells ablation (Figure S1A). Interestingly, layer-specific changes in the different types of spines emerged, with a decrease in thin spine density in SR and SO, but an increase in the number of mushroom spines in SO (Figure S1A).

**Figure 2.**
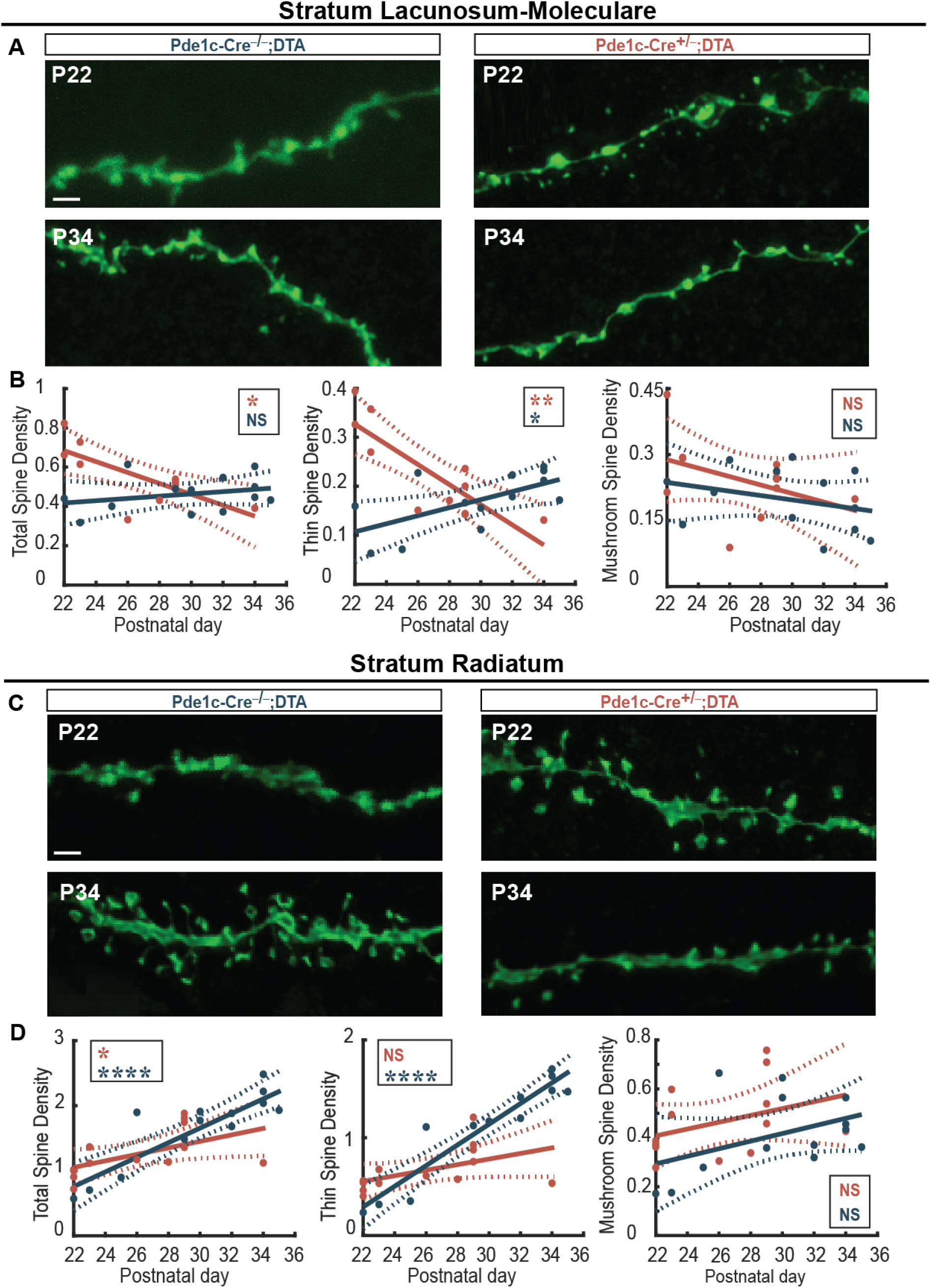
Ablation of CR cells disrupts the formation of dendritic spines on CA1 pyramidal cells. (A) Images of terminal dendritic segments in SLM of CA1 pyramidal cells in control (n=13, from 8 animals) and Pde1c-Cre^+/-^ ;DTA mice (n=13, from 6 animals). (B) Simple regression models of total (*left*: Pde1c-Cre^+/-^;DTA: R^2^=0.54 vs. Pde1c-Cre^-/-^;DTA: R^2^=0.08), thin (*middle*: R^2^=0.7, vs Pde1c-Cre^-/-^;DTA:R^2^=0.38) and mushroom (*right*: Pde1c-Cre^+/-^ ;DTA: R^2^=0.17), Pde1c-Cre^-/-^;DTA: R^2^= 0.09) spine densities in SLM across developmental time points in control and Pde1c-Cre^+/-^;DTA. (C) Images of terminal dendritic segments in SR of CA1 pyramidal cells in control and Pde1c-Cre^+/-^;DTA mice. (D) Simple regression models of total (*left*: *R*^*2*^=0.77 vs. *R*^*2*^=.32), thin (*middle*: *R*^*2*^=0.89 vs. *R*^*2*^=0.25) and mushroom (*right*: R^2^=0.18 vs. R^2^ = 0.13) spine densities in SR across developmental time points in control and Pde1c-Cre^+/-^;DTA. Scale bars in (A) and (C) 1 µm, density levels in (B) and (D) measured as number of spines per 1 µm. p-values: NS ≥ 0.05, * ≤ 0.05, ** ≤ 0.001, **** ≤ 0.0001.

Spine formation happens over several weeks, with most immature synapses starting to acquire more mature properties from the first postnatal week and onward (Basarsky et al., 1994; Holtmaat et al., 2005). To assess if the developmental stage influenced the number of spines, we applied a simple linear regression model to correlate postnatal age with spine density. Dendritic spines of the SLM demonstrated opposite trends as total spine density is reduced with age in Pde1c-Cre^+/-^ ;DTA but does not change in the Pde1c-Cre^-/-^;DTA (Figure 2B), with thin spines demonstrating the most drastic decrease (Figure 2B). Mushroom (Figure 2B), stubby and filopodia (data not shown) were non-significantly changed in both experimental groups.

In SR and SO, the total spine density is significantly different between the two groups, with the Pde1c-Cre^-/-^;DTA group showing a clear increase, that we could not observe in the Pde1c-Cre^+/-^;DTA mice (Figures 2D, S1D). As in the SLM, this effect is likely due to significant changes in thin spine densities (Figure 2D, S1D). No differences were observed in mushroom (Figure 2D, S1D), stubby and filopodia spines (data not shown).

Previous reports show that as CA1 pyramidal neurons mature, the density of dendritic spines doubles, and that the number of the different types of spines changes non-uniformly (Harris et al., 1992; Holtmaat et al., 2005). Indeed, our control mice show an increase in spine density as postnatal development proceeds, with different magnitudes across layers. However, in the absence of CR cells, spine development is severely impaired. The observed layer-specific differences suggests that distinct inputs to CA1 pyramidal cells are differentially affected by CR cells ablation.

### CR cells-dependent changes of gene regulatory networks

To understand molecular underpinnings of the spine changes observed in the CR cells ablated hippocampus, dorsal hippocampi were microdissected from Pde1c-Cre^-/-^;DTA and Pde1c-Cre^+/-^;DTA groups at P15 and P30. Total RNA was extracted from dissected hippocampal tissue and mRNA was sequenced (Figure 3A). Fastq reads were mapped to the Mmv79 genome map and its annotated transcriptome using Kallisto and resultant mapped reads per gene were normalized using the TMM method in R (Figure S2A). Individual samples were then clustered via PCA analysis and grouped by study design (Figure 3B). PC1 and PC2 were driven by age-related differences between the P15 and P30 groups whereas PC2 and PC3 were more related to the injection. Each group clustered with respect to each other revealing differences between P15 groups though P30 Pde1c-Cre^+/-^;DTA was more variable in its clustering and thus did not show separation from Pde1c-Cre^-/-^;DTA group. Next, we investigated differential gene expression between groups using a contrast matrix. A linear model was then established using the voom variance trend (Figure S2B) and the normalized gene expression to analyze contrast matrices of all possible comparisons between genotypic, age, and treatment groups. A decrease in CR cell-specific markers confirmed CR cells deletion (Figure S2D). Differentially expressed genes were analyzed by establishing volcano plots of ΔP15 (Figure S2E), ΔP30 (Figure S2F), ΔCre^-^ (Figure S2G), and ΔCre^+^ (Figure 3C) groups. No genes were found to be significantly changed in either the ΔP15 (Figure S2E) or ΔP30 (Figure S2F), except for pan neuronal-associated gene, *Atxn7l3b*. As observed in Figure 3C, significant changes in gene expression were observed in the ΔCre^-^ (Figure S2G) and ΔCre^+^ (Figure 3D) groups further demonstrating that molecular changes mediated by CR cells may occur subtly over this developmental period.

**Figure 3.**
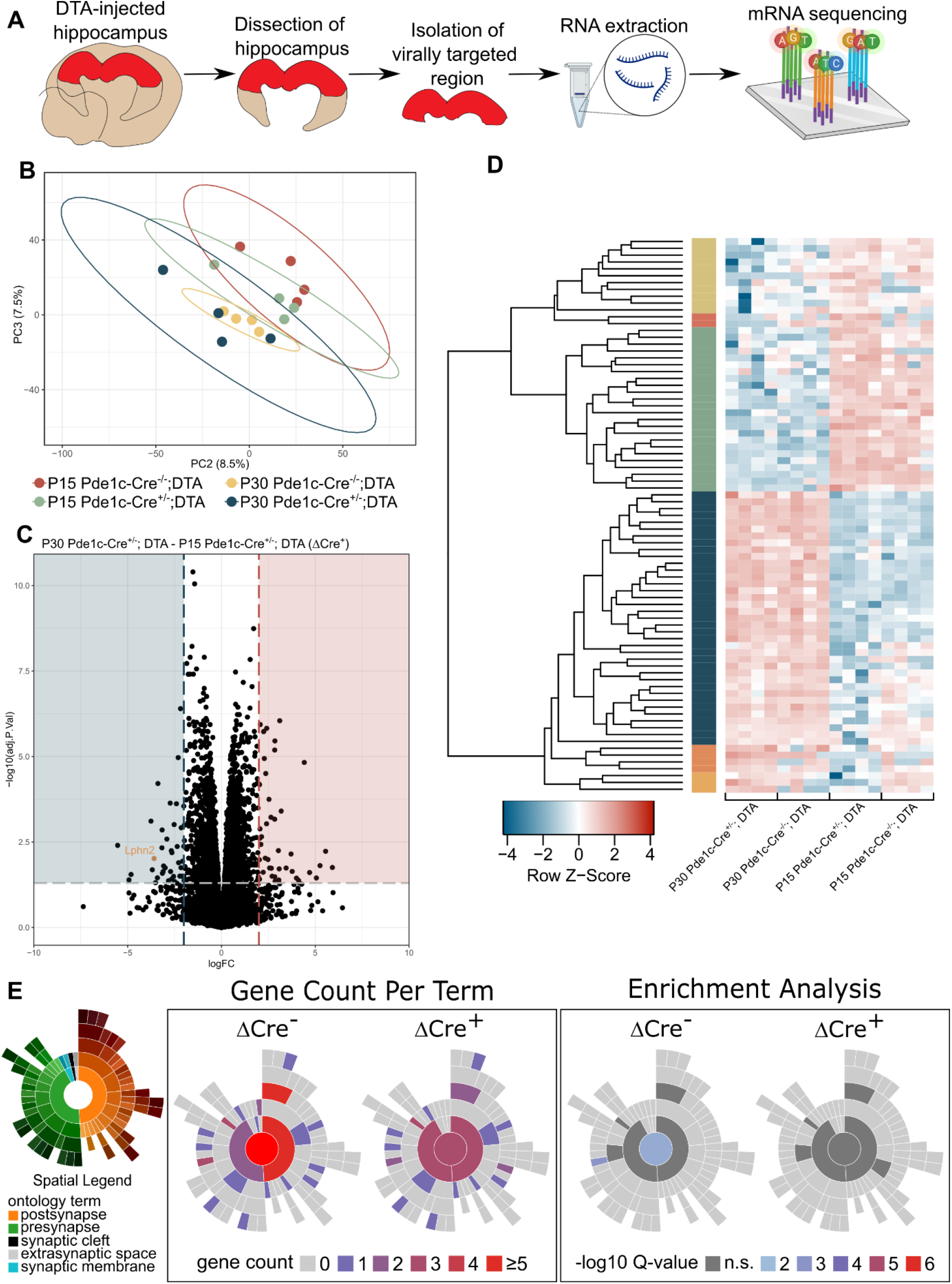
CR cells-dependent changes of gene regulatory networks. (A) Schematic of mRNA sequencing of injected hippocampi from P15 Pde1c-Cre^-/-^;DTA (n=4), P15 Pde1c-Cre^+/-^;DTA (n=4), P30 Pde1c-Cre^-/-^;DTA (n=4), and P30 Pde1c-Cre^+/-^;DTA (n=4). (B) PCA of each group plotted as the second and third principal components as related to injection. (C) Volcano plot of ΔCre^+^ group, demonstrating significant transcriptional changes in 94 genes. Colored vignettes indicate threshold values at LogFC≥2 and adj p-val≤0.05. Values in blue are downregulated over time whereas values in red are upregulated over time in the CR cell ablated group. (D) Heatmap clustering analysis of individual gene expression of individual samples between groups. Genes shown were thresholded at LogFC≥2 and adj p-val≤0.05. Genetic and sample clustering were performed using Kendall’s correlation method. Colored modules between branches and heatmap cells indicate clustering groups (k=6) driving differences between groups. From the top, the second, fifth, and sixth modules indicate genetic expression differences with respect to CR cells deletion. (E) SynGO analysis of top 200 genes sorted by adjusted p-val in the ΔCre groups demonstrating loss of overall gene terms in cellular compartment as well as significance in enrichment analysis. Spatial legend indicates corresponding location on plots, gene count indicates total term number enriched in each cellular compartment, and -log10Q-value indicates Benjamini-Hochberg corrected p-val.

To understand how individual gene expressions contribute to the uniqueness of each study design group’s transcriptional profile, a heatmap analysis (Figure 3D) was made of the same gene lists comprising the volcano plots (Figures S2E-G, 3C). Resultant modules included 3 gene clusters with noticeable difference between injection groups (Clusters 2, 5, and 6) and one gene cluster of subtle changes between groups (Cluster 1). Cluster 2 included *Recql4* and *Socs1*, cluster 5 included *Gm14440, Kansl2, Tcte1*, and *Tgtp2*, and Cluster 6 included *Sh3tc2, Fam166b*, and *Hmga1*. Genes like *Socs1* (Baker et al., 2009) and *Tgtp2* (Meijer et al., 2022), encode for proteins known to be involved in CNS immunoresponse and are activated by interferon pathways. Genes involved in neuronal development including neuronal migration like *Tcte1* (Zur Lage et al., 2019), chromatin remodeling complex genes required for neurogenesis like *Recql4* (Kohzaki et al., 2021), *Kansl2* (Ferreyra Solari et al., 2016), and *Hmga1* (Vignali & Marracci, 2020), or myelination genes like *Sh3tc2* (Arnaud et al., 2009) were significantly changed between injection groups in both timepoints (Figure 3D). Additionally, neuronal outgrowth-related gene, *Fam166b* (Varga et al., 1995) was downregulated in the P15 Pde1c-Cre^+/-^;DTA group.

Based on these expression changes a ranked gene list gene ontology analysis was performed by sorting genes by descending Log_2_FC. Neural-related terms were plotted using enrichR to visualize individual enriched genes and their distribution with respect to the P15 or P30 in the ΔCre^-^ and ΔCre^+^ groups (Figure S3A). This analysis revealed conserved ontologies between the developmental groups, like ensheathment of neurons and hormone activity. However, several ontological terms were not present after CR cells ablation including neuron recognition, neuron projection guidance, and neuropeptide signaling pathways. As anticipated, these pathways were enriched in the P15 Pde1c-Cre^-/-^ group, suggesting that CR cells ablation might affect axonal pathfinding. Overall, these results show that the absence of CR cells alters the developmental trajectory of the postnatal hippocampus and gene regulatory networks centered around neurogenic chromatin remodeling complexes and projection growth-related genes.

To next investigate the resultant synaptic changes in neurodevelopmental genes and gene networks, a second analysis was performed focused on synapse-related ontologies. Ranked gene lists were established for ΔCre^-^ and ΔCre^+^ groups and were analyzed using SynGO (Koopmans et al., 2019). SynGO analysis returned a significant enrichment in synaptic cellular compartment ontology overall in the ΔCre^-^ group but a loss in both absolute term number and significance in the ΔCre^+^ group (Figure 3E). Likewise, synaptic term loss and insignificant enrichment in the ΔCre^+^ group was conserved within the biological processes term analysis (Figure S3B). Analysis of individual terms from cellular compartment gene ontology revealed a loss in term number and significance in presynaptic and postsynaptic terms but most notable loss of enrichment occurred at the presynaptic level in cytosolic- and membrane-related terms (Figure S3C). Similarly, biological process gene ontology analysis revealed a loss in term number and significance in synaptic processes including synaptic organization and modulation of chemical synaptic transmission (Figure S3D). Additionally, a complete loss of terms in the ΔCre^+^ group was observed for synapse assembly, process in the postsynapse, and neurotransmitter localization to postsynaptic membrane. These findings reflect the loss of spines observed in the morphological analysis and further suggest that CR cells mediate connectivity in the postnatal developing hippocampus. Overall, these findings reinforce CR cells’ role in facilitating synaptic establishment in the developing hippocampus.

### SLM specific effect of CR-cells ablation

Bulk RNAseq analysis of the whole hippocampus revealed significant downregulation of synaptic mRNA after postnatal ablation of CR cells. A deeper analysis of the significantly downregulated genes in the ΔCre^+^ group showed that a unique synaptic marker, *Lphn2*, downregulated in the ΔCre^-^ group but was ∼8 times more downregulated in the ΔCre^+^ group (Figure 3C, yellow text). *Lphn2* encodes for the postsynaptic guidance molecule, Latrophilin-2. In dendritic layers of the hippocampus, latrophilin-2 is exclusively expressed in the CA1 SLM and acts as an axonal guidance molecule for inputs from the entorhinal cortex (Anderson et al., 2017; Donohue et al., 2021; Pederick et al., 2021). To understand if *Lphn2* is topographically downregulated, CA1 SLM was microdissected from Pde1c-Cre^-/-^;DTA and Pde1c-Cre^+/-^;DTA groups at P15 and P30. Total RNA was extracted from the tissue and qRT-PCR analysis was performed targeting genes specific for CA1 synapses (*Lphn2*, (Donohue et al., 2021) *Robo2* (Blockus et al., 2021) or CA3 synapses (*Igsf8, Neo1* (Apóstolo et al., 2020)) to assess specificity of changes (Figure 4A). qRT-PCR analysis revealed that *Lphn2* and *Robo2* were both significantly upregulated at P15 Pde1c-Cre^+/-^;DTA compared to Pde1c-Cre^-/-^;DTA and conversely, were significantly downregulated at P30 Pde1c-Cre^-/-^;DTA compared to Pde1c-Cre^-/-^;DTA (Figure 4B). No changes were observed in *Igsf8* and *Neo1* expression. The deletion of CR cells inverts the expression trend of synaptic markers responsible for unique inputs to CA1 and may explain the trends observed in spine density (Figure 2). Immunohistochemistry confirmed a reduction in *Lphn2* in the SLM at P30 in CR cells-ablated mice (Figure 4C). Latrophilin-2 is thus downregulated both at the protein and mRNA levels and further demonstrates that CR cells may have a lasting role in synapse maturation.

**Figure 4.**
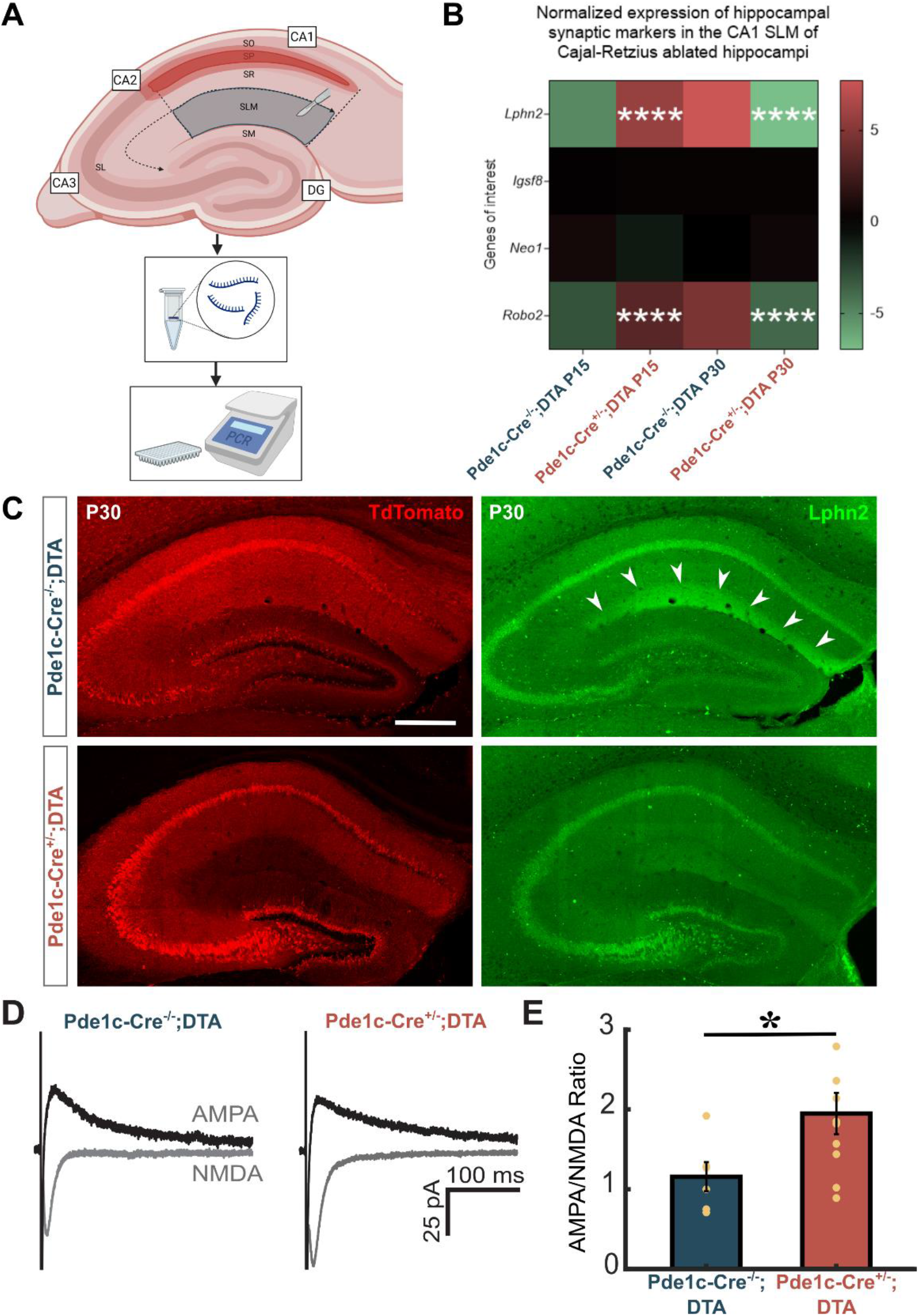
SLM-specific effect of CR cells ablation. (A) Schematic of microdissection of CA1 SLM from hippocampi of P15 Pde1c-Cre^-/-^;DTA (n=5), P15 Pde1c-Cre^+/-^;DTA (n=6), P30 Pde1c-Cre^-/-^;DTA (n=2), and P30 Pde1c-Cre^+/-^ ;DTA (n=5). (B) Heatmap depicting log2FC of qRT-PCR-acquired expression of *Adgrl2 (Lphn2), Igsf8, Neo1*, and *Robo2*. (C) Image of coronal section depicting Lphn2 expression in Pde1c-Cre^-/-^;DTA and Pde1c-Cre^+/-^;DTA groups at P30. (D) Representative sample traces of evoked EPSCAMPA and EPSCNMDA recorded in the same cell at -70 mV and +40 mV, respectively, from a Pde1c-cre^-/-^;DTA (n=8) and Pde1c^+/-^;DTA (n=10). (E) Summary graph for AMPA/NMDA ratio. Scale bar in (C) is 300 µm. p-values: *<0.05., ****<0.0001

To investigate whether the changes observed in spines and synaptic-related genes in SLM are reflected at the physiological level we determined the AMPA/NMDA ratio of evoked excitatory synaptic currents (eEPSCs) recorded in CA1 pyramidal neurons upon stimulation of fibers in SLM (Figure 4D). The ratio was significantly higher in the Pde1c-Cre^+/-^;DTA group compared to Pde1c-Cre^-/-^;DTA (Figure 4E). Since thalamic fibers localized in SLM seems to exclusively target GABAergic interneurons (Andrianova et al., 2021), our data suggest drastic changes in the entorhinal-CA1 connectivity.

In summary, our findings reveal that CR cells appear to be crucial for circuitry establishment in the mouse hippocampus during postnatal development.

## Discussion

CR-cells are widely understood to have a critical role in cortical lamination, through the release of reelin (D’Arcangelo et al., 1995; Ma et al., 2014). Their disappearance at early stages of postnatal development in the neocortex (Ma et al., 2014) had often led to neglecting their postnatal role. However, in the last years, studies focused on interventions targeting CR-cell densities in the neocortex have shown effects on spine density, dendritic complexity, excitatory/inhibitory balance and thalamic input integration (de Frutos et al., 2016; Genescu et al., 2022; Riva et al., 2019). These studies greatly highlighted the importance of CR cells during the development of neocortical regions. What role CR cells play in the hippocampus, where they persist for several months (Anstötz et al., 2016; Ma et al., 2014; Supèr et al., 1998), however, remains a crucial unanswered question.

In this study we generated a model to ablate CR cells from the hippocampus specifically at postnatal stages. Postnatal ablation of hippocampal CR cells led to 1) layer-specific alterations in the number and type of dendritic spines on CA1 pyramidal cells; 2) changes in the developmental trajectory of the postnatal hippocampus; 3) downregulation in the expression of synaptic-related genes; 4) significant reduction in the mRNA and protein levels of Lhpn2, a gene important for the establishment and maintenance of the entorhinal-hippocampal circuit.

By inducing early CR cell death, we were able to reveal a critical role of CR cells in the postnatal hippocampus. What mechanisms lead to the observed changes in the hippocampal circuits, however, are yet to be determined. Reelin, the main protein correlated to CR cells’ function, has been shown to influence synaptogenesis of hippocampal neurons (Bosch et al., 2016; Niu et al., 2008). However, CR cells also release glutamate, contacting both GABAergic cells and CA1 pyramidal neurons (Quattrocolo & Maccaferri, 2014) and, according to the synaptotrophic hypothesis, glutamatergic synaptic transmission regulates synaptogenesis (Cline & Haas, 2008). As our sequencing analysis did not reveal drastic changes in any one specific pathway, we expect the observed phenotype to be related to changes in several convergent pathways modulated by CR cells. Future work focused on more refined manipulation will likely reveal insights in the mechanism underlying the specific alterations we reported.

Pharmacological ablation of CR cells in organotypic slice cultures prevents the ingrowth of entorhinal cortex fibers into the hippocampus (Del Río et al., 1997). Since, the axon of CR cells extends to several areas of the hippocampal region, reaching the entorhinal cortex (Anstötz et al., 2016; Deng et al., 2007; Supèr et al., 1998), it has been proposed that fibers from the entorhinal cortex could use CR cells axons as a scaffold to reach the hippocampus, where they will transiently contact CR cells (Supèr et al., 1998). Here, we can contribute to the proposed role of CR cells as regulators of the development of the entorhinal-hippocampal circuit. In fact, the absence of hippocampal CR cells resulted in a reduction of Lphn2. Lphn2 has been proposed to function as a postsynaptic guidance molecule for the hippocampal network, with peak expression at P8 during development (Pederick et al., 2021), and as an adhesion molecule for the maintenance of these connections in the adult (Anderson et al., 2017). The observed changes on Lphn2 suggest that CR cells contribute to the maturation of the entorhinal-hippocampal circuit also during postnatal development. What genes/proteins are implicated in the disruption of the other subcircuits of the hippocampus causing our layer-specific changes are yet to be determined.

The hippocampal formation matures during the first weeks of postnatal development (Alberini & Travaglia, 2017; Donato et al., 2017; Dumas, 2005), as hippocampal function slowly emerges (Wills et al., 2013). In the future, it will be important to address how the observed changes affect the function of the different areas of the hippocampal formation.

In conclusion, our work reveals a crucial role for CR cells in the maturation of the hippocampal circuit, highlighting the importance of this transient cell population in the region and opening new and interesting questions on their impact on hippocampal function.

## Acknowledgements

We would like to thank B.A. Zaharia for excellent technical assistance; R.N. Raveendran and the Viral Vector Core Facility of the Kavli Institute for Systems Neuroscience (NTNU) for viral reagents; H.M. Møllergård for genotyping; P.J.B. Girão for assistance with microscopy; G. Fishell, E. Moser, M. Witter and all the members of the Quattrocolo lab for their insightful comments throughout this project. We further thank the staff in the animal facility at the Kavli Institute for Systems Neuroscience. Bulk RNA sequencing, adaptor trimming, and quality control was provided by the Genomics Core Facility (GCF), Norwegian University of Science and Technology (NTNU). GCF is funded by the Faculty of Medicine and Health Sciences at NTNU and Central Norway Regional Health Authority. The work was supported by a Research Council of Norway (RCN) FRIPRO grants to G.Q. (grant number 324305), the European Union’s Horizon 2020 Research and Innovation Programme (Marie Skłodowska-Curie grant agreement to G.Q.: CaRe-Space no. 839988), the Trond Mohn Foundation (Mohn Research Center of the Brain, grant number 2021TMT04, to G.Q.), an RCN Centre of Excellence grant (Centre of Neural Computation, grant number 223262; NORBRAIN, grant number 295721, to G.Q.), and the Kavli Foundation (to G.Q.).

## Author contribution

Conceptualization, I.L.G., K.D. and G.Q.; supervision, B.v.L. and G.Q.; experiment design, I.L.G., K.D., N.P.M., B.v.L., P.S. and G.Q.; data collection, I.L.G., K.D., N.P.M., M.J.N., H.K. and G.Q.; visualization, I.L.G. and K.D.; writing, original draft, I.L.G., K.D. and G.Q.; writing, review, and editing, I.L.G., K.D., N.P.M., M.J.N., B.v.L., P.S. and G.Q.; funding acquisition: B.v.L. and G.Q.; discussion, comments, all of the authors.

## Experimental procedures

### Animals

Male mice belonging to the Pde1c-Cre transgenic mouse line (B6.FVB(Cg)-Tg(Pde1c-cre)IT146Gsat/Mmucd, MMRRC 030708) (Osheroff & Hatten, 2009) were bred with wild type females (C57BL/6JBomTac, Taconic). Both male and female mice originating from this cross were used for experiments in this study.

All experiments were conducted in compliance with protocols approved by the Norwegian Food Safety Authorities and European Directive 2010/63/EU (FOTS ID 24847). All mice were housed in enriched environment cages in a 12 hr light/dark cycle with food and water ad libitum. Pups were separated from the mother at postnatal day 21.

### Viral Injection Procedure

All pups were subjected to bilateral injections with a recombinant adeno-associated virus (pAAV-mCherry-flex-dtA Addgene plasmid #58536; http://n2t.net/addgene:58536 ; RRID:Addgene_58536) (Wu et al., 2014) expressing Cre-dependent diphtheria toxin A fragment (AAV-DTA) at postnatal day zero (P0). All animals were anesthetized with isoflurane (3%) and head fixed in a stereotaxic frame (Koppf) with custom-made adaptor. The skin was stretched using standard lab tape with a diamond cut. Injection coordinates were then calculated from *lambda* (AP: +0.8; ML: ±1.2; Z: −1.22) in each mouse. A virus-filled glass pipette (Drummond) was attached to an injector, Nanoject III (Drummond Scientific Company), used for the injections (48.4 nl, divided in 4 injections of 12.1 volume, 0.10 rate, with 2 seconds delay).

Injected Cre^+^ animals (Pde1c-Cre^+/-^;DTA) were considered the experimental group and their Cre^-^ littermates (Pde1c-Cre^-/-^;DTA) were considered controls. Only animals which had cells expressing mCherry in the outer blade of the dentate gyrus were used for experiments as this indicated viral spread to most of the hippocampus.

### Tissue Preparation

All animals were first anesthetized with isoflurane before being euthanized with a lethal intraperitoneal injection of pentobarbital (100 mg/Kg). They were then transcardially perfused first with phosphate buffer saline (PBS) followed by a 4% paraformaldehyde (PFA) solution in PBS. Brains were harvested and postfixed 3 hours in PFA. They were then moved to a 15% sucrose solution overnight (stored at 4°C) and then to a 30% sucrose solution for an additional overnight. Fixed brains were coronally sectioned on a freezing microtome (ThermoScientific) at a 50µm thickness and stored in cryoprotective solution (40% PBS, 30% glycerol, 30% ethylene glycol).

### Immunohistochemistry

For immunohistochemistry, sections were rinsed in PBS for 10 minutes and subsequentially blocked for 30 minutes in a Blocking Solution containing 10% Normal Donkey Serum (Jackson ImmunoResearch,) and 0.1% Triton X-100 (Merk) dissolved in PBS. Antigen retrieval was performed once, only for staining of Latrophilin-2, when sections were heated in a basic antigen retrieval reagent (Sodium Citrate, pH 8.5) at 75 C for 10 minutes before blocking. Sections were then incubated with primary antibodies diluted in an Incubating Solution containing 1% Normal Donkey Serum and 0.1% Triton X-100 dissolved in PBS, for three overnights in 4 C with gentle shaking. Primary antibodies were as follows: rat anti-RFP, dilution 1:1000 (Chromotek, #5F8); rabbit anti-p73, dilution 1:500 (Abcam, #ab40658); mouse anti-reelin (Abcam, #ab78540), dilution 1:500; rabbit anti-Lphn2, dilution 1:1000 (Novus Biologicals, NBP2-58704). Preparation of the secondary antibody staining included 3×1 hour rinsing with PBS which was followed by overnight incubation in secondary antibodies (biotinylated IgG, 1:500 diluted in Incubating Solution) in 4°C. Secondary antibodies included: donkey anti-mouse Alexa 488; donkey anti-rabbit Alexa 488; donkey anti-rat Alexa 594; donkey anti-rabbit Alexa 647. The next day, sections were rinsed 3×10 minutes in PBS, then incubated in PBS with 1:200 Nissl (Invitrogen, N21479) for 1 hour. Sections were then rinsed in PBS-0.1% Tx for 10 minutes followed by one-hour rinse in PBS. Lastly, sections were mounted on Super-frost slides (Thermo Fisher Scientific) in a petri dish with PBS, briefly rinsed with dH_2_O, cover slipped with mounting medium (Fluoromount-G, ThermoFisher), then sealed with nail polish and stored in 4°C.

### Confocal Imaging and Cell Counts

A LSM880 Zeiss Confocal Microscope (Zeiss) was used to acquire tiled z-stack images of the hippocampal fissure and molecular layer of the dentate gyrus using a Zeiss plan apochromat 20x/numerical aperture 0.8 objective. Four coronal sections from each animal were chosen for imaging. Sections were 150 μm apart demonstrating a minimum viral spread of 600 μm.

Confocal imaging files containing the dorsal hippocampus were uploaded in Neurolucida (Micro Bright Field Bioscience) for analysis. A contour delineating the stratum lacunosum-moleculare and the molecular layer of the inner blade of the dentate gyrus was created. Symbols for each signal was used to count cells labeled by different markers. To quantify double or triple labeled cells, a different marker was used for each combination of signals. We counted all cells contained within the contours. Since non-consecutive slices were counted, overcounting the z-axis was not relevant, and no correction was applied. Quantification of the number of markers in the contour was done in Neurolucida Explorer (Micro Bright Field Bioscience) and exported in excel files. To assess the specificity of the Pde1c-cre mouse line, the number of cells was used to calculate the ratio (Figure1B). To assess the effect of the AAV-DTA virus, the same delineations were made but only cells immunolabeled with p73 were counted. The density of neurons labeled by a marker was measured as the number of labeled neurons in a contour divided by the area included in the contour (cells/mm^2^).

### Slice Preparation and Electrophysiological Recordings

Acute brain slices for recordings were prepared from P22–P35 Pde1c-Cre mice, previously injected at P0 with AAV-mCherry-flex-dtA. Mice were anesthetized with Isufluorane and injected I.P. with pentobarbital after sedation. Transcardial perfusion was performed with chilled modified artificial cerebrospinal fluid (ACSF, in mM): NMDG 93, KCl 3, NaH_2_PO_4_ 1.25, NaHCO_3_ 25, Hepes 20, Glucose 10, NAC 10, MgCl_2_ 5, CaCl_2_ 0.5, saturated with 95% O_2_, 5% CO_2_ (pH 7.4). The brain was removed and placed into a small container filled with chilled modified ACSF. Coronal sections (300 µm) were cut using a vibrating microtome (VT1000, Leica) and slices were incubated at 34 C-35 C for at least 30 min and then stored at RT until use in a modified ACSF (in mM): NaCl 124, KCl, 3, NaH_2_PO_4_ 1.25, NaHCO_3_ 30, Hepes 20, Glucose 10, NAC 10, MgCl_2_ 5, CaCl_2_ 0.5, saturated with 95% O_2_, 5% CO_2_ (pH 7.4). Slices were transferred to an upright microscope (BW51, Olympus). Slices were perfused with preheated ACSF of the following composition (in mM): NaCl 124, KCl 3, NaH_2_PO_4_ 1.25, NaHCO_3_ 26, Glucose 10, MgCl_2_ 1, CaCl_2_ 1.6 saturated with 95% O2, 5%CO2 (pH 7.4) and maintained at a constant temperature (30 C–32 C). 10 µM Gabazine was added to recording solution before starting of experiments. Recoding pipettes were pulled from borosilicate glass capillaries (Sutter Instruments) and had a resistance of 3-5 MΩ when filled with the internal solution of the following composition: Cs-MeSo_3_ 130, CsCl 5, HEPES 10, EGTA 0.2, Mg-ATP 4, Na-GTP 0.3, Phosphocreatine-Na_2_ 5, QX-314-Cl 5 (pH 7.3). Biocytin was added each day of recordings to reach a 0.3%-0.5% final concentration. Whole-cell recordings were performed using a Multiclamp 700B amplifier (Molecular Devices) with Digidata 1550A and a computer with pClamp 11 (Molecular Devices). Voltage-clamp signals were filtered at 4 kHz and recorded with a sampling rate of 10 kHz. Recordings were performed at a holding potential of -70 mV and +40 mV. No correction was made for the junction potential between the pipette and the ACSF. Fluorescence from the injected virus was checked for each slice, and recordings were performed only if signal was detected in both Stratum Pyramidale of CA1 and the Granular Layer of the Dentate Gyrus (inner blade). Neurons located in Stratum Pyramidale of CA1 hippocampus were randomly selected. The following number of cells were recorded in each condition: Pde1c-Cre^-/-^;DTA: 8 cells from 7 animals; Pde1c-Cre^+/-^;DTA: 10 cells from 6 animals. Fluorescence expression and targeting of the viral injection was confirmed post-hoc during recovery of the morphology of pyramidal cells. Cells with a non-pyramidal neuron morphology were excluded from all the analyses.

Postsynaptic currents were evoked by placing a borosilicate glass filled with recording solution in stratum lacunosum-moleculare. Positive current was generated through an ISO-Flex stimulator and intensity was adjusted to reliably evoke a consistent response of small amplitude. After recording AMPA-mediated EPSCs at -70 mV, the recording solution containing 10 µM DNQX (6,7-dinitroquinoxaline-2,3-dione) was washed in the recording chamber. Once the amplitude of the EPSC_AMPA_ was reduced to approximately zero, the holding potential of the cell was depolarized to +40 mV and NMDA-mediated EPSCs were recorded.

### Dendritic Reconstruction and Spine Counting

Cells were filled with biocytin during whole-cell recording and then fixed overnight in 4% paraformaldehyde in 0.1 M PBS at 4 C. Fluorescent and biocytin signals were detected with the following procedure. Sections were rinsed in PBS 3 times, 5-10 minutes for each rinse, and then blocked for an hour in Blocking Solution with 0.5% Triton. Slices were then moved in Incubating Solution with 0.5% Triton with Rat anti-RFP antibody (1:500, Chromoteck) for 3 overnights at 4 C with gentle shaking. Slices were then rinsed 3 times for an hour in PBS and then moved into Incubating Solution with 0.5% Triton with Streptavidin-conjugated Alexa Fluor 488 (1:500, Invitrogen) and Donkey anti-Rat secondary antibodies conjugated with Alexa Fluor 594 (1:500, Invitrogen) and left incubating overnight at 4 C with gentle shaking. The following day, slices were rinsed 5 times for 10 minutes in PBS and then mounted onto SuperFrost glass slides (Thermo Fisher Scientific). Neurons were imaged using a LSM880 Zeiss Confocal Microscope (Zeiss) with a Zeiss plan apochromat 63x/numerical aperture 1.4 oil immersion objective. Each neuron was imaged as part of a three-fold experiment: imaging three dendritic segments in the SLM, the SR and lastly the SO. Dendritic segments were chosen based on the same criteria: each segment is part of a terminal dendrite defined as the last bifurcation of the dendritic shaft. Cells without apical dendrites reaching SLM were excluded from the experiment. This resulted in a distribution of n cells per postnatal day: P22 (n_Pde1c-Cre-/-;DTA_=1; n_Pde1c-Cre+/- ;DTA_=4), P23 (n_Pde1c-Cre-/-;DTA_ =1; n_Pde1c-Cre+/-;DTA_ =2), P25 (n_Pde1c-Cre-/-;DTA_ =1), P26 (n_Pde1c-Cre-/-;DTA_ =1, n_Pde1c-Cre+/- ;DTA_ =1), P28 (n_Pde1c-Cre+/-;DTA_ =1), P29 (n_Pde1c-Cre-/-;DTA_ =1, n_Pde1c-Cre+/-;DTA_ =4), P30 (n_Pde1c-Cre-/-;DTA_ =2), P32 (n_Pde1c-Cre-/-;DTA_ =2), P33 (n_Pde1c-Cre-/-;DTA_ =2), P34 (n_Pde1c-Cre-/-;DTA_ =3, n_Pde1c-Cre+/-;DTA_ =1), P35 (n_Pde1c-Cre-/-;DTA_ =1). All data acquisition and analysis were performed by an experimenter blinded to the mouse genotype.

To reverse the chromatic aberration by the confocal microscope, the image was deconvoluted by Huygens Deconvolution Software (Essential pack, Scientific Volume Imaging) using a Classic Maximum Likelihood Estimation protocol with a theoretical point spread function (PSF) generated by the Huygens software based on the properties of the immersion objective. Dendritic shafts were reconstructed on a Neurolucida Software (Micro Bright Field Bioscience) by means of user-guided tracing with directional kernels. Spines were tagged using unique markers to classify spine subtypes. This spine classification was based on established criteria’s (Harris et al., 1992). Spines were subsequently quantified by means of Branch Structure Analysis and Markers and Regions Analysis on Neurolucida Explorer (Micro Bright Field Bioscience).

### Whole hippocampus extraction

P30 Pde1c-Cre mice, previously injected at P0 with AAV-mCherry-flex-DTA were first anesthetized with isoflurane before being euthanized with a lethal intraperitoneal injection of pentobarbital (100 mg/Kg). After decapitation, brain was removed from the skull and immersed in an ice-cold buffer (320 mM sucrose, 10 mM Hepes, ph=7.4). The hippocampi were then dissected and checked for injection reporter using a Stereo Microscope Fluorescent Adapter with a Green lamp (Nightsea). Samples were included only if an intense fluorescence was detected in the dorsal hippocampus and only the fluorescent part was dissected.

### Microdissection of CA1 SLM

P15 and P30 Pde1c-Cre mice, previously injected at P0 with AAV-mCherry-flex-DTA, were first anesthetized with isoflurane before being euthanized with a lethal intraperitoneal injection of pentobarbital (100 mg/Kg). After decapitation, brain was removed from the skull and immersed in chilled sucrose ACSF with the following composition: NaCl 87, KCl 2.5, NaH_2_PO_4_ 1.25, NaHCO_3_ 26, Sucrose 75, Glucose 10, MgCl_2_ 7, CaCl_2_ 0.5, saturated with 95% O_2_, 5% CO_2_ (pH 7.4). Coronal sections (500 µm) were cut using a vibrating microtome (VT1000S, Leica). Slices containing the dorsal hippocampus were quickly moved to a petri dish containing chilled sucrose ACSF and correct targeting of the viral injection was assessed with the use of a Stereo Microscope Fluorescent Adapter with a Green lamp (Nightsea). Only slices in which bright fluorescence was visible in the entire hippocampus were used for the microdissection. A V8 Discovery Stereoscope (Zeiss) was used to guide the microdissection of the CA1 SLM. The CA1 SLM was finitely dissected using a Barkan microknife (FST) and Dumont #5 forceps (FST). Tissue was taken from two subsequent slice sections from both hemispheres and was kept on ice until RNA extraction.

### RNA Extraction

Whole hippocampus or microdissected CA1 SLM was processed for RNA extraction. Tissue was homogenized and processed using either Qiagen RNAEasy (Qiagen) or Nucleospin RNA extraction kit (Machery-Nagel). Isolated RNA from whole hippocampus was analyzed using bulk mRNA Seq whereas isolated RNA from CA1 SLM was analyzed using qRT-PCR.

### Bulk mRNA Sequencing

RNA from Snap frozen injected whole hippocampus tissue was quality controlled using Agilent 2100 Bioanalyzer and RNA quantity was detected by Qubit fluorometric quantitation. Samples were sequenced at a minimum RIN value of 8.0. Libraries were performed using the Illumina® Stranded mRNA Prep, Ligation kit. High throughput sequencing was performed using an NS500HO flowcell (Illumina) for an average read of ∼16×10^6^ reads. Fastq files were assessed using bcl2fastq/QC report. Sequencing was performed at the Genomics Core Facility at NTNU.

### Transcriptome Analysis

Raw read files were assessed for quality and then pseudoaligned raw reads to GRCm39 *Mus musculus* using kallisto (Bray et al., 2016). Aligned transcriptomic data processing and analysis was executed in RStudio (RStudio IDE 2021.09.0 Build 351) using tximport (Soneson et al., 2015), ensembldb (Rainer et al., 2019), edgeR (Robinson et al., 2010), matrixStats, limma (Ritchie et al., 2015), GSEABase (Morgan et al., 2022), GSVA (Hänzelmann et al., 2013), ClusterProfiler (Koopmans et al., 2019), enrichplot (Yu, 2022), and SynGO (Koopmans et al., 2019). Raw reads were normalized using trimmed means of M-values (TMM) and genes were disregarded if the expression <1 in at least 4 samples. Significant gene expression was determined using threshold values of LogFC ≥2 and Benjamini-Hochberg adjusted p-val ≤ 0.05. SynGO analysis was performed using the top 200 genes (<1% of kept genes) in a ranked list of adjusted p-values. All samples were analyzed from P15 Pde1c-Cre^-/-^;DTA (n=4), P15 Pde1c-Cre^+/-^;DTA (n=4), P30 Pde1c-Cre^-/-^;DTA (n=4), P30 Pde1c-Cre^+/-^;DTA (n=4), P15 Pde1c-Cre^-/-^;Control (n=2), P15 Pde1c-Cre^+/-^;Control (n=2), P30 Pde1c-Cre^-/-^;Control (n=2), P30 Pde1c-Cre^+/-^;Control (n=2) to control for genotype variability at each age until pairwise analyses were performed. Plots were made using cowplot and ggplot.qRT-PCR

RNA from microdissected CA1 SLM samples were processed using Reverse Transcriptase Core Kit 300 (Eurogentec RT-RTCK-03). Approximately 100 ng of RNA was reverse transcribed into cDNA for qRT-PCR analysis. SensiFAST SYBR mix (12 μl, BioLine BIO-98020) and cDNA (8 μl) were combined in tubes and quantified using StepOne*Plus* (Thermo Fisher). Primers were designed using NIH Primer Blast with the most recent genome sequencing of *Mus musculus*. All primers were designed with predicted amplicon product ranges of 50-150 bp. Take-off cycles thresholds (C_T_) for each gene were normalized to their sample’s respective *Gapdh* take off-cycle.

### Primers used for each gene

*Gapdh* Forward: CTGCACCACCAACTGCTTAG; *Gapdh* Reverse: ATCCACAGTCTTCTGGGTGG; *Lphn2* Forward: TCTGGAACAGAGGGCGAAG; *Lphn2* Reverse: TAGCCAGAAGGAATATGGAAGAGT; *Igsf8* Forward: CAGGAGTCGCCCTAGTTACC; *Igsf8* Reverse: TGGCTGGAAGACAGTCAACA; *Neo1* Forward: AAAGGCCTCCCGAAAAAGTG; *Neo1* Reverse: AGCAGCGACTCGGAAGTTAT; *Robo2* Forward: TCATGGGTCCCAAAACTTGC; *Robo2* Reverse: TGCAGGCACTGATGGTAGAA.

### Statistical Analysis

For cell, spine, and electrophysiological AMPA/NMDA current ratio quantification MATLAB_R2019b (The MathWorks Inc) was used to extract all statistical measures. Number of cells per µm^2^ area was converted to mm^2^ and then averaged across four sections of each brain. Brain averages of CR cells densities per mm^2^ were statistically compared through a two-sample t-test assuming unequal variance.

Raw dendritic spine data was standardized by dendritic length to correct for variations in dendritic branching and uneven sample numbers between groups. As such, spine density for each dendritic segment was given by the spine quantity divided by branch length (µm). This equation was applied to the total spine density (comprising all spine markers) and subcategories which included unique markers for mushroom, thin, stubby and filopodia spines. A simple linear regression model was applied to quantify age-related effects in the data across all hippocampal layers using postnatal age as the predictor variable.

To calculate the AMPA/NMDA current ratio we first averaged the EPSC_AMPA_ and EPSC_NMDA_ recorded events and calculated the peak amplitude. Those values were used then to calculate the ratio. These current ratios were then statistically compared by means of a two-sample t-test.

Statistical analysis of qRT-PCR data was performed using Graphpad Prism v8.0.2. DCt values were compared between groups for each gene using mixed-effects multiple comparison analysis corrected using FDR in two-stage step-up method of Benhamini, Krieger, and Yekutieli approach with a Q threshold of 1%. This analysis was chosen to adjust for number of samples, number of genes, and number of groups analyzed. Statistical significance is reported as the adjusted p-value.

### GEO Accession

Bulk RNAseq data from whole hippocampus samples are registered under: [GEO name] [link].

## Supplementary Figures

**Supplementary Figure 1.**
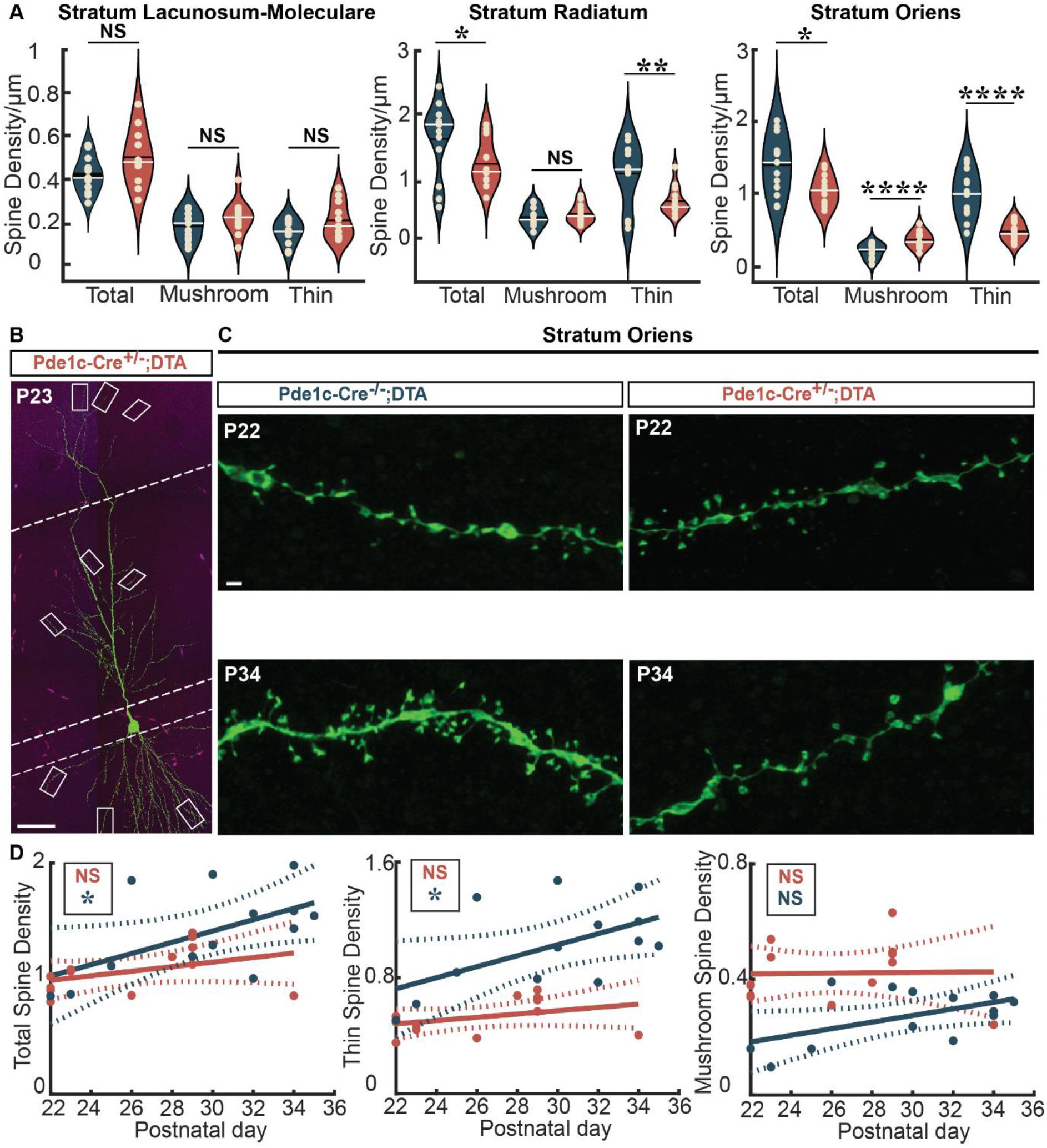
Layer specificity in dendritic spine development. (A) Total, mushroom and thin spine densities per µm measured in SLM (Pde1c-Cre^-/-^;DTA mean = 0.47; Pde1c-Cre^+/-^;DTA mean = 0.55), SR (mean = 1.65, mean=1.23) and SO (mean = 1.40, mean = 1.06) of of Pde1c-Cre^-/-^;DTA (*blue*) and of Pde1c-Cre^+/-^;DTA (*red*). Black line signifies mean, white line median. (B) Example pyramidal cell demonstrating dendrite selection. Each selected dendrite was terminal in the layer of interest (e.g., taken after the last bifurcation). Delineations showing SLM (*top*), SR (*middle*), SO (*bottom*). (C) Examples of Pde1c-Cre^-/-^;DTA and Pde1c-Cre^+/-^;DTA terminal dendritic segments in SO at P22 and P35. (D) Simple regression models of Pde1c-Cre^-/-^;DTA and Pde1c-Cre^+/-^;DTA total (*left*: R^2^=0.32 vs. R^2^=0.15), thin (*middle*: R^2^=0.31 vs. R^2^=0.14) and mushroom (*right*: R^2^ >0.01 vs. R^2^=0.141) spine densities in SO across developmental time points. Scale bar in (B) 50 µm and in (C) 1 µm. p-values: NS ≥ 0.05, * ≤ 0.05, ** ≤ 0.001, **** ≤ 0.0001.

**Supplementary Figure 2.**
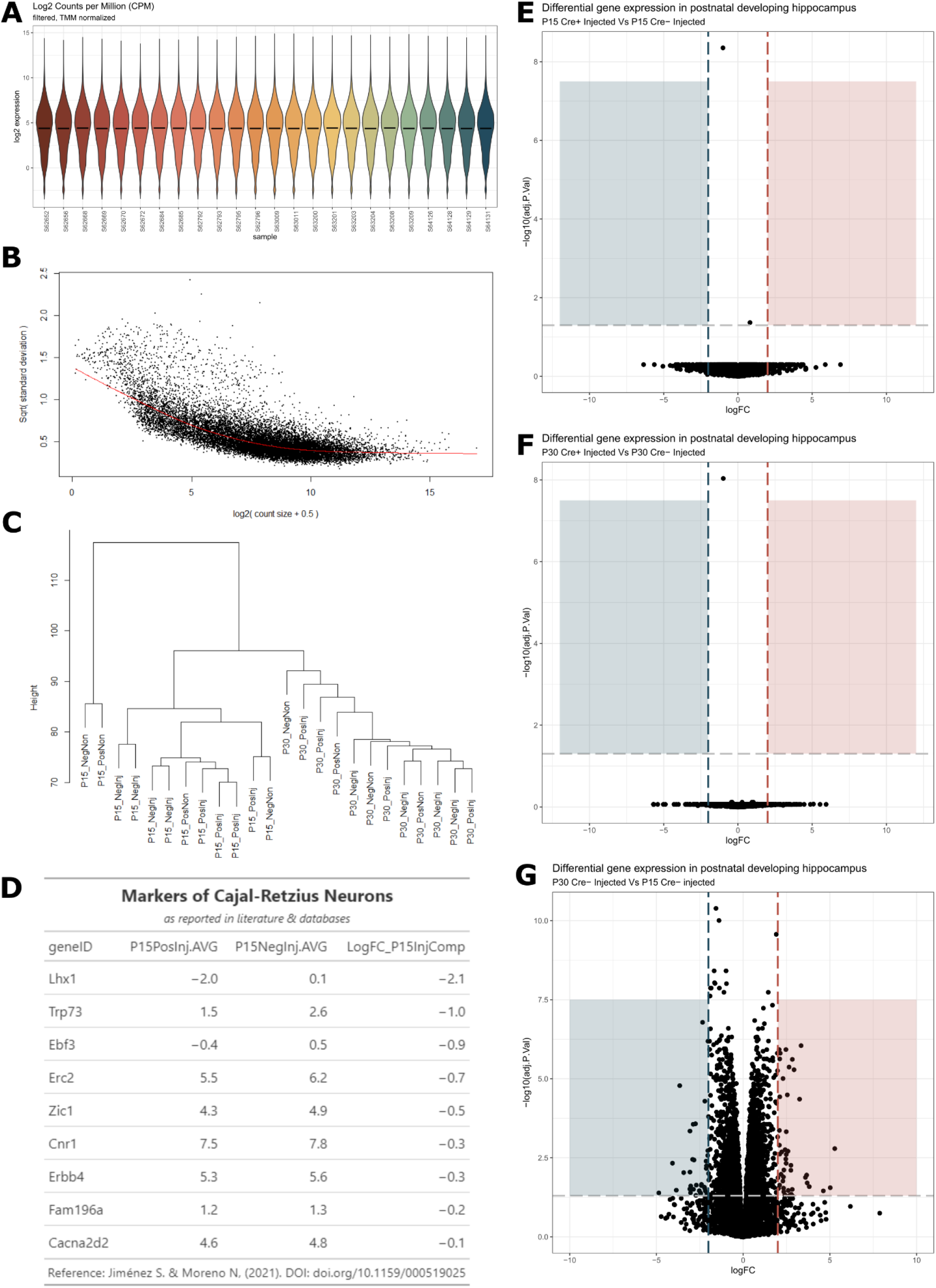
Transcriptomic analysis of injected hippocampi of Pde1c-Cre^-/-^ and Pde1c-Cre^+/-^ at P15 and P30. (A) Trimmed-means of M-values of 14,555 genes and their log2FC of expression in samples from groups (n=24) including: P15 Pde1c-Cre^-/-^;DTA (n=4), P15 Pde1c-Cre^+/-^;DTA (n=4), P30 Pde1c-Cre^-/-^;DTA (n=4), P30 Pde1c-Cre^+/-^;DTA (n=4), P15 Pde1c-Cre^-/-^;Control (n=2), P15 Pde1c-Cre^+/-^;Control (n=2), P30 Pde1c-Cre^-/-^;Control (n=2), P30 Pde1c-Cre^+/-^;Control (n=2). (B) Voom means variance plot of individual genes (black points) across all samples. Red slope indicates fit used for Bayesian analysis of contrast matrices in limma. (C) Sample clustering using Manhattan method for distance and average agglomeration method for height mapping. (D) Table of commonly acknowledged CR cells markers extracted from literature including *Lhx1, Trp73, Ebf3, Erc2, Zic1, Cnr1, Erbb4, Fam196a*, and *Cacna2d2* (Jiménez & Moreno, 2021). Log2FC differences generated by the following equation: [*Gene*]_P15 Pde1c-Cre+/-;DTA_ – [*Gene*]_P15 Pde1c-Cre-/-;DTA_. (E) Volcano plot of changes in gene expression at P15 between injected groups. (F) Volcano plot of changes in gene expression at P30 between injected groups. Colored vignettes indicate threshold values at LogFC ≥2 and adj p-val≤0.05. (G) Volcano plot of ΔCre^-^ group, demonstrating significant transcriptional changes in ∼70 genes. In F-G, Colored vignettes indicate threshold values at LogFC ≥2 and adj p-val≤0.05. Values in blue are significantly downregulated over time whereas values in red are significantly upregulated.

**Supplementary Figure 3.**
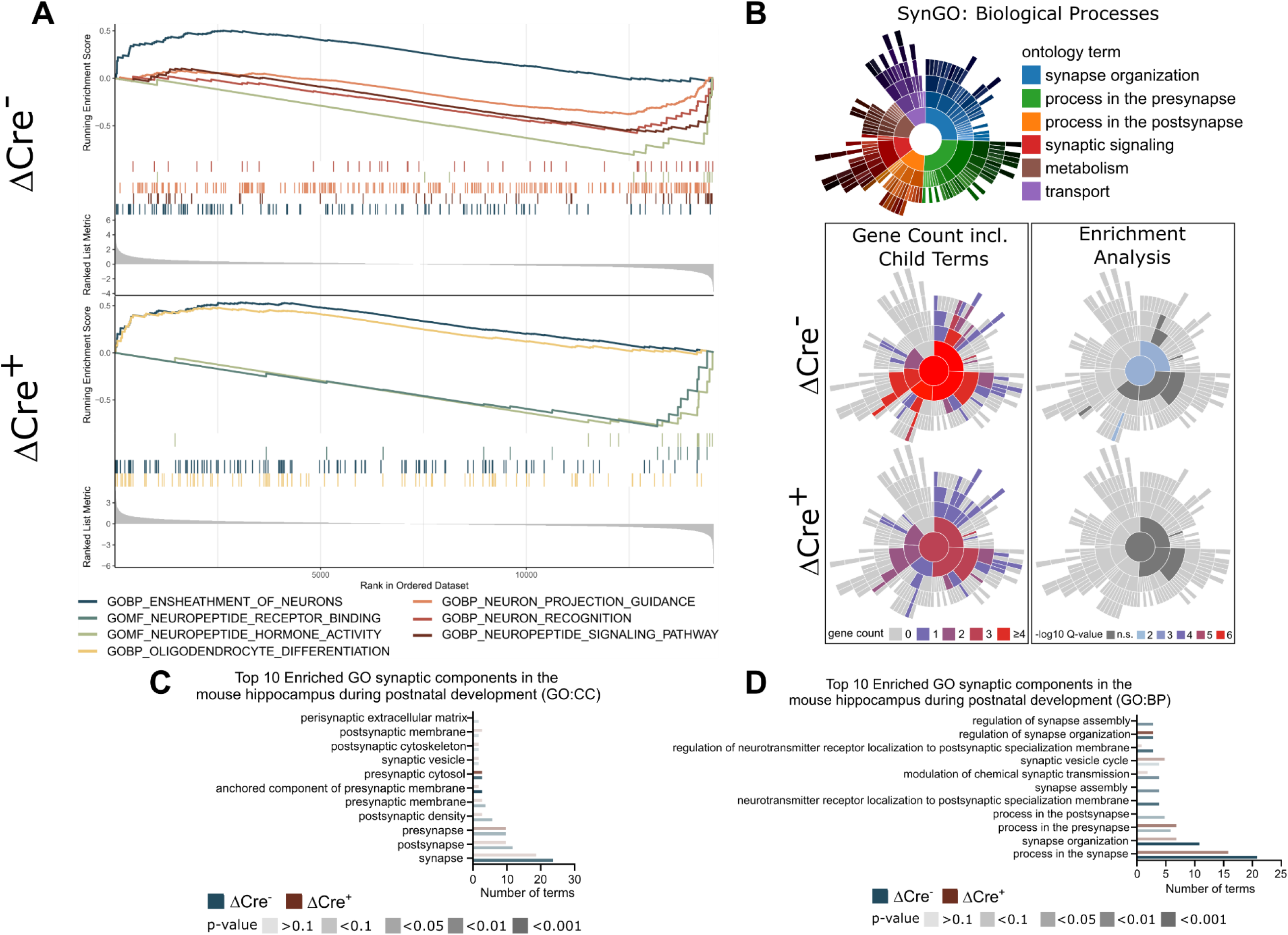
Gene ontology analysis of injected hippocampi of Pde1c-Cre^-/-^ and Pde1c-Cre^+/-^ at P15 and P30. (A) Enrichr plots generated between ΔCre^-^ and ΔCre^+^ groups highlighting differences in select developmental gene networks. Plots generated by LogFC-ranked gene list of ∼14,555 genes in each group and indicate changes from GSEA’s GO database in either biological processes (GOBP) or molecular function (GOMF). Ensheathment of neurons and hormone activity are conserved between both groups while the remaining 5 ontological annotations are unique to either. (B) SynGO analysis of top 200 genes sorted by adjusted p-val in the ΔCre groups demonstrating loss of overall gene terms in biological processes as well as significance in enrichment analysis. Spatial legend indicates corresponding process on plots, gene count indicates total term number enriched in each cellular compartment, and -log10Q-value indicates Benjamini-Hochberg corrected p-val. (C) Top 10 enriched GO CC synaptic terms between ΔCre^-^ and ΔCre^+^ groups from SynGO analysis. ΔCre^-^ values are depicted in blue and ΔCre^+^ values are depicted in red. Adj. p-values are depicted by saturation of colors. Complimentary to Figure 3E. (D) Top 10 enriched GO BP synaptic terms between ΔCre^-^ and ΔCre^+^ groups from SynGO analysis. ΔCre^-^ values are depicted in blue and ΔCre^+^ values are depicted in red. Adj. p-values are depicted by saturation of colors. Complimentary to Supplementary Figure 3B.

| REAGENT/RESOURCE | SOURCE | RRID |
| --- | --- | --- |
| **Antibodies** |  |  |
| Rat anti-RFP | Chromotek | RRID:AB_2336064 |
| Rabbit anti-p73 | Abcam | RRID:AB_776999 |
| Mouse anti-reelin | Abcam | RRID:AB_1603148 |
| Rabbit anti-Lphn2 | Novus Biologicals | Not registered |
| **Plasmid** |  |  |
| pAAV-mCherry-flex-dta | UNC vector core, Boston, USA | RRID:Addgene_58536 |
| **Virus** |  |  |
| AAV-mCherry-flex-dta | UNC vector core, Boston, USA |  |
| AAV-mCherry-flex-dta | Kavli Vector Core, Trondheim, Norway | Not registered |
| **Software and algorithms** |  |  |
| Neurolucida 360° | MBF Bioscience | RRID:SCR_016788 |
| Huygens Software | Essential pack Scientific Volume Imaging | RRID:SCR_014237 |
| MATLAB | The MathWorks Inc | RRID:SCR_001622 |
| Black Zen Software | Carl Zeiss Microscopy | RRID:SCR_018163 |
| Adobe Illustrator | Adobe Inc | RRID:SCR_010279 |
| Adobe Photoshop | Adobe | RRID:SCR_014199 |
| R Project for Statistical Computing | Open | RRID:SCR_001905 |
| **Experimental models:**<br>**Organisms/strains** |  |  |
| Pde1c-Cre<br>IT146Gsat/Mmucd | MMRRC | RRID:MMRRC_030708-UCD |
| C57BL/6JBomTac | Taconic | RRID:IMSR_TAC:b6jbom |
| Ai9 (B6.Cg-Gt(ROSA)26Sortm9(CAG-tdTomato)Hze/J) | The Jackson Laboratory | RRID:IMSR_JAX:007909 |

## References

Alberini, C. M., & Travaglia, A. (2017). Infantile Amnesia: A Critical Period of Learning to Learn and Remember. J Neurosci, 37(24), 5783–5795. https://doi.org/10.1523/jneurosci.0324-17.2017

Anderson, G. R., Maxeiner, S., Sando, R., Tsetsenis, T., Malenka, R. C., & Südhof, T. C. (2017). Postsynaptic adhesion GPCR latrophilin-2 mediates target recognition in entorhinal-hippocampal synapse assembly. The Journal of cell biology, 216(11), 3831–3846. https://doi.org/10.1083/jcb.201703042

Andrianova, L., Brady, E. S., Margetts-Smith, G., Kohli, S., McBain, C. J., & Craig, M. T. (2021). Hippocampal CA1 pyramidal cells do not receive monosynaptic input from thalamic nucleus reuniens. bioRxiv, 2021.2009.2030.462517. https://doi.org/10.1101/2021.09.30.462517

Anstötz, M., Huang, H., Marchionni, I., Haumann, I., Maccaferri, G., & Lübke, J. H. (2016). Developmental Profile, Morphology, and Synaptic Connectivity of Cajal-Retzius Cells in the Postnatal Mouse Hippocampus. Cereb Cortex, 26(2), 855–872. https://doi.org/10.1093/cercor/bhv271

Anstötz, M., Quattrocolo, G., & Maccaferri, G. (2018). Cajal-Retzius cells and GABAergic interneurons of the developing hippocampus: Close electrophysiological encounters of the third kind. Brain Res, 1697, 124–133. https://doi.org/10.1016/j.brainres.2018.07.028

Apóstolo, N., Smukowski, S. N., Vanderlinden, J., Condomitti, G., Rybakin, V., Ten Bos, J., Trobiani, L., Portegies, S., Vennekens, K. M., Gounko, N. V., Comoletti, D., Wierda, K. D., Savas, J. N., & de Wit, J. (2020). Synapse type-specific proteomic dissection identifies IgSF8 as a hippocampal CA3 microcircuit organizer. Nat Commun, 11(1), 5171. https://doi.org/10.1038/s41467-020-18956-x

Arnaud, E., Zenker, J., de Preux Charles, A.S., Stendel, C., Roos, A., Médard, J. J., Tricaud, N., Kleine, H., Luscher, B., Weis, J., Suter, U., Senderek, J., & Chrast, R. (2009). SH3TC2/KIAA1985 protein is required for proper myelination and the integrity of the node of Ranvier in the peripheral nervous system. Proc Natl Acad Sci U S A, 106(41), 17528–17533. https://doi.org/10.1073/pnas.0905523106

Baker, B. J., Akhtar, L. N., & Benveniste, E. N. (2009). SOCS1 and SOCS3 in the control of CNS immunity. Trends Immunol, 30(8), 392–400. https://doi.org/10.1016/j.it.2009.07.001

Basarsky, T. A., Parpura, V., & Haydon, P. G. (1994). Hippocampal synaptogenesis in cell culture: developmental time course of synapse formation, calcium influx, and synaptic protein distribution. J Neurosci, 14(11 Pt 1), 6402-6411. https://doi.org/10.1523/jneurosci.14-11-06402.1994

Blockus, H., Rolotti, S. V., Szoboszlay, M., Peze-Heidsieck, E., Ming, T., Schroeder, A., Apostolo, N., Vennekens, K. M., Katsamba, P. S., Bahna, F., Mannepalli, S., Ahlsen, G., Honig, B., Shapiro, L., de Wit, J., Losonczy, A., & Polleux, F. (2021). Synaptogenic activity of the axon guidance molecule Robo2 underlies hippocampal circuit function. Cell Rep, 37(3), 109828. https://doi.org/10.1016/j.celrep.2021.109828

Bosch, C., Muhaisen, A., Pujadas, L., Soriano, E., & Martínez, A. (2016). Reelin Exerts Structural, Biochemical and Transcriptional Regulation Over Presynaptic and Postsynaptic Elements in the Adult Hippocampus. Front Cell Neurosci, 10, 138. https://doi.org/10.3389/fncel.2016.00138

Bray, N. L., Pimentel, H., Melsted, P., & Pachter, L. (2016). Near-optimal probabilistic RNA-seq quantification (vol 34, pg 525, 2016). Nature Biotechnology, 34(8), 888–888. https://doi.org/DOI10.1038/nbt0816-888d

Buzsáki, G., & Moser, E. I. (2013). Memory, navigation and theta rhythm in the hippocampal-entorhinal system. Nat Neurosci, 16(2), 130–138. https://doi.org/10.1038/nn.3304

Ceranik, K., Deng, J., Heimrich, B., Lübke, J., Zhao, S., Förster, E., & Frotscher, M. (1999). Hippocampal Cajal-Retzius cells project to the entorhinal cortex: retrograde tracing and intracellular labelling studies. Eur J Neurosci, 11(12), 4278–4290. https://doi.org/10.1046/j.1460-9568.1999.00860.x

Cline, H., & Haas, K. (2008). The regulation of dendritic arbor development and plasticity by glutamatergic synaptic input: a review of the synaptotrophic hypothesis. J Physiol, 586(6), 1509–1517. https://doi.org/10.1113/jphysiol.2007.150029

D’Arcangelo, G., G. Miao, G., Chen, S.-C., Scares, H. D., Morgan, J. I., & Curran, T. (1995). A protein related to extracellular matrix proteins deleted in the mouse mutant reeler. Nature, 374(6524), 719–723. https://doi.org/10.1038/374719a0

de Frutos, Cristina A., Bouvier, G., Arai, Y., Thion Morgane S., Lokmane, L., Keita, M., Garcia-Dominguez, M., Charnay, P., Hirata, T., Riethmacher, D., Grove Elizabeth A., Tissir, F., Casado, M., Pierani, A., & Garel, S. (2016). Reallocation of Olfactory Cajal-Retzius Cells Shapes Neocortex Architecture. Neuron, 92(2), 435–448. https://doi.org/https://doi.org/10.1016/j.neuron.2016.09.020

Del Río, J. A., Heimrich, B., Borrell, V., Förster, E., Drakew, A., Alcántara, S., Nakajima, K., Miyata, T., Ogawa, M., Mikoshiba, K., Derer, P., Frotscher, M., & Soriano, E. (1997). A role for Cajal– Retzius cells and reelin in the development of hippocampal connections. Nature, 385(6611), 70–74. https://doi.org/10.1038/385070a0

Deng, J. B., Yu, D. M., Wu, P., & Li, M. S. (2007). The tracing study of developing entorhino-hippocampal pathway. Int J Dev Neurosci, 25(4), 251–258. https://doi.org/10.1016/j.ijdevneu.2007.03.002

Donato, F., Jacobsen, R. I., Moser, M. B., & Moser, E. I. (2017). Stellate cells drive maturation of the entorhinal-hippocampal circuit. Science, 355(6330). https://doi.org/10.1126/science.aai8178

Donohue, J. D., Amidon, R. F., Murphy, T. R., Wong, A. J., Liu, E. D., Saab, L., King, A. J., Pae, H., Ajayi, M. T., & Anderson, G. R. (2021). Parahippocampal latrophilin-2 (ADGRL2) expression controls topographical presubiculum to entorhinal cortex circuit connectivity. Cell Reports, 37(8), 110031. https://doi.org/https://doi.org/10.1016/j.celrep.2021.110031

Dumas, T. C. (2005). Late postnatal maturation of excitatory synaptic transmission permits adult-like expression of hippocampal-dependent behaviors. Hippocampus, 15(5), 562–578. https://doi.org/10.1002/hipo.20077

Ferreyra Solari, N. E., Belforte, F. S., Canedo, L., Videla-Richardson, G. A., Espinosa, J. M., Rossi, M., Serna, E., Riudavets, M. A., Martinetto, H., Sevlever, G., & Perez-Castro, C. (2016). The NSL Chromatin-Modifying Complex Subunit KANSL2 Regulates Cancer Stem-like Properties in Glioblastoma That Contribute to Tumorigenesis. Cancer Res, 76(18), 5383–5394. https://doi.org/10.1158/0008-5472.Can-15-3159

Genescu, I., Aníbal-Martínez, M., Kouskoff, V., Chenouard, N., Mailhes-Hamon, C., Cartonnet, H., Lokmane, L., Rijli, F. M., López-Bendito, G., Gambino, F., & Garel, S. (2022). Dynamic interplay between thalamic activity and Cajal-Retzius cells regulates the wiring of cortical layer 1. Cell Reports, 39(2), 110667. https://doi.org/https://doi.org/10.1016/j.celrep.2022.110667

Gil, V., Nocentini, S., & Del Río, J. A. (2014). Historical first descriptions of Cajal-Retzius cells: from pioneer studies to current knowledge. Frontiers in neuroanatomy, 8, 32–32. https://doi.org/10.3389/fnana.2014.00032

Hänzelmann, S., Castelo, R., & Guinney, J. (2013). GSVA: gene set variation analysis for microarray and RNA-Seq data. BMC Bioinformatics, 14(1), 7. https://doi.org/10.1186/1471-2105-14-7

Harris, K. M., Jensen, F. E., & Tsao, B. (1992). Three-dimensional structure of dendritic spines and synapses in rat hippocampus (CA1) at postnatal day 15 and adult ages: implications for the maturation of synaptic physiology and long-term potentiation. J Neurosci, 12(7), 2685–2705. https://doi.org/10.1523/jneurosci.12-07-02685.1992

Holtmaat, A. J., Trachtenberg, J. T., Wilbrecht, L., Shepherd, G. M., Zhang, X., Knott, G. W., & Svoboda, K. (2005). Transient and persistent dendritic spines in the neocortex in vivo. Neuron, 45(2), 279–291. https://doi.org/10.1016/j.neuron.2005.01.003

Jiménez, S., & Moreno, N. (2021). Analysis of the Expression Pattern of Cajal-Retzius Cell Markers in the Xenopus laevis Forebrain. Brain Behav Evol, 1–20. https://doi.org/10.1159/000519025

Kohzaki, M., Ootsuyama, A., Umata, T., & Okazaki, R. (2021). Comparison of the fertility of tumor suppressor gene-deficient C57BL/6 mouse strains reveals stable reproductive aging and novel pleiotropic gene. Scientific Reports, 11(1), 12357. https://doi.org/10.1038/s41598-021-91342-9

Koopmans, F., van Nierop, P., Andres-Alonso, M., Byrnes, A., Cijsouw, T., Coba, M. P., Cornelisse, L. N., Farrell, R. J., Goldschmidt, H. L., Howrigan, D. P., Hussain, N. K., Imig, C., de Jong, A. P. H., Jung, H., Kohansalnodehi, M., Kramarz, B., Lipstein, N., Lovering, R. C., MacGillavry, H., Mariano, V., Mi, H., Ninov, M., Osumi-Sutherland, D., Pielot, R., Smalla, K. H., Tang, H., Tashman, K., Toonen, R. F. G., Verpelli, C., Reig-Viader, R., Watanabe, K., van Weering, J., Achsel, T., Ashrafi, G., Asi, N., Brown, T. C., De Camilli, P., Feuermann, M., Foulger, R. E., Gaudet, P., Joglekar, A., Kanellopoulos, A., Malenka, R., Nicoll, R. A., Pulido, C., de Juan-Sanz, J., Sheng, M., Südhof, T. C., Tilgner, H. U., Bagni, C., Bayés, À., Biederer, T., Brose, N., Chua, J. J. E., Dieterich, D. C., Gundelfinger, E. D., Hoogenraad, C., Huganir, R. L., Jahn, R., Kaeser, P. S., Kim, E., Kreutz, M. R., McPherson, P. S., Neale, B. M., O’Connor, V., Posthuma, D., Ryan, T. A., Sala, C., Feng, G., Hyman, S. E., Thomas, P. D., Smit, A. B., & Verhage, M. (2019). SynGO: An Evidence-Based, Expert-Curated Knowledge Base for the Synapse. Neuron, 103(2), 217-234.e214. https://doi.org/10.1016/j.neuron.2019.05.002

Ma, J., Yao, X. H., Fu, Y., & Yu, Y. C. (2014). Development of layer 1 neurons in the mouse neocortex. Cereb Cortex, 24(10), 2604–2618. https://doi.org/10.1093/cercor/bht114

Meijer, M., Agirre, E., Kabbe, M., van Tuijn, C. A., Heskol, A., Zheng, C., Mendanha Falcão, A., Bartosovic, M., Kirby, L., Calini, D., Johnson, M. R., Corces, M. R., Montine, T. J., Chen, X., Chang, H. Y., Malhotra, D., & Castelo-Branco, G. (2022). Epigenomic priming of immune genes implicates oligodendroglia in multiple sclerosis susceptibility. Neuron, 110(7), 1193-1210.e1113. https://doi.org/https://doi.org/10.1016/j.neuron.2021.12.034

Meyer, G., Socorro, A. C., Garcia, C. G. P., Millan, L. M., Walker, N., & Caput, D. (2004). Developmental Roles of p73 in Cajal-Retzius Cells and Cortical Patterning. The Journal of Neuroscience, 24(44), 9878. https://doi.org/10.1523/JNEUROSCI.3060-04.2004

Morgan, M., Falcon, S., & Gentleman, R. (2022). GSEABase: Gene set enrichment data structures and methods. In (Version R package version 1.58.0.)

Niu, S., Yabut, O., & D’Arcangelo, G. (2008). The Reelin signaling pathway promotes dendritic spine development in hippocampal neurons. The Journal of neuroscience : the official journal of the Society for Neuroscience, 28(41), 10339–10348. https://doi.org/10.1523/JNEUROSCI.1917-08.2008

Osheroff, H., & Hatten, M. E. (2009). Gene Expression Profiling of Preplate Neurons Destined for the Subplate: Genes Involved in Transcription, Axon Extension, Neurotransmitter Regulation, Steroid Hormone Signaling, and Neuronal Survival. Cerebral Cortex, 19(Suppl_1), i126–i134. https://doi.org/10.1093/cercor/bhp034

Pederick, D. T., Lui, J. H., Gingrich, E. C., Xu, C., Wagner, M. J., Liu, Y., He, Z., Quake, S. R., & Luo, L. (2021). Reciprocal repulsions instruct the precise assembly of parallel hippocampal networks. Science, 372(6546), 1068–1073. https://doi.org/10.1126/science.abg1774

Quattrocolo, G., & Maccaferri, G. (2013). Novel GABAergic circuits mediating excitation/inhibition of Cajal-Retzius cells in the developing hippocampus. J Neurosci, 33(13), 5486–5498. https://doi.org/10.1523/jneurosci.5680-12.2013

Quattrocolo, G., & Maccaferri, G. (2014). Optogenetic activation of cajal-retzius cells reveals their glutamatergic output and a novel feedforward circuit in the developing mouse hippocampus. The Journal of neuroscience : the official journal of the Society for Neuroscience, 34(39), 13018–13032. https://doi.org/10.1523/JNEUROSCI.1407-14.2014

Rainer, J., Gatto, L., & Weichenberger, C. X. (2019). ensembldb: an R package to create and use Ensembl-based annotation resources. Bioinformatics, 35(17), 3151–3153. https://doi.org/10.1093/bioinformatics/btz031

Ramón y Cajal, S., Sols Lucia, A., & Reinoso Suárez, F. (2011). Recuerdos de mi vida: historia de mi labor científica.

Ritchie, M. E., Phipson, B., Wu, D., Hu, Y., Law, C. W., Shi, W., & Smyth, G. K. (2015). limma powers differential expression analyses for RNA-sequencing and microarray studies. Nucleic Acids Res, 43(7), e47. https://doi.org/10.1093/nar/gkv007

Riva, M., Genescu, I., Habermacher, C., Orduz, D., Ledonne, F., Rijli, F. M., Lopez-Bendito, G., Coppola, E., Garel, S., Angulo, M. C., & Pierani, A. (2019). Activity-dependent death of transient Cajal-Retzius neurons is required for functional cortical wiring. Elife, 8. https://doi.org/10.7554/eLife.50503

Robinson, M. D., McCarthy, D. J., & Smyth, G. K. (2010). edgeR: a Bioconductor package for differential expression analysis of digital gene expression data. Bioinformatics, 26(1), 139–140. https://doi.org/10.1093/bioinformatics/btp616

Soneson, C., Love, M. I., & Robinson, M. D. (2015). Differential analyses for RNA-seq: transcript-level estimates improve gene-level inferences. F1000Res, 4, 1521. https://doi.org/10.12688/f1000research.7563.2

Supèr, H., Martinéz, A., Del Rió, J.A., & Soriano, E. (1998). Involvement of Distinct Pioneer Neurons in the Formation of Layer-Specific Connections in the Hippocampus. The Journal of Neuroscience, 18(12), 4616. https://doi.org/10.1523/JNEUROSCI.18-12-04616.1998

Travaglia, A., Bisaz, R., Sweet, E. S., Blitzer, R. D., & Alberini, C. M. (2016). Infantile amnesia reflects a developmental critical period for hippocampal learning. Nature Neuroscience, 19(9), 1225–1233. https://doi.org/10.1038/nn.4348

Varga, Z. M., Schwab, M. E., & Nicholls, J. G. (1995). Myelin-associated neurite growth-inhibitory proteins and suppression of regeneration of immature mammalian spinal cord in culture. Proceedings of the National Academy of Sciences of the United States of America, 92(24), 10959–10963. https://doi.org/10.1073/pnas.92.24.10959

Vignali, R., & Marracci, S. (2020). HMGA Genes and Proteins in Development and Evolution. Int J Mol Sci, 21(2). https://doi.org/10.3390/ijms21020654

Wills, T. J., Muessig, L., & Cacucci, F. (2013). The development of spatial behaviour and the hippocampal neural representation of space. Philosophical transactions of the Royal Society of London. Series B, Biological sciences, 369(1635), 20130409–20130409. https://doi.org/10.1098/rstb.2013.0409

Wu, Z., Autry, A. E., Bergan, J. F., Watabe-Uchida, M., & Dulac, C. G. (2014). Galanin neurons in the medial preoptic area govern parental behaviour. Nature, 509(7500), 325–330. https://doi.org/10.1038/nature13307

Yu, G. (2022). enrichplot: Visualization of Functional Enrichment Result. In (Version R package version 1.16.0)

Yuste, R., Hawrylycz, M., Aalling, N., Aguilar-Valles, A., Arendt, D., Armañanzas, R., Ascoli, G. A., Bielza, C., Bokharaie, V., Bergmann, T. B., Bystron, I., Capogna, M., Chang, Y., Clemens, A., de Kock, C. P. J., DeFelipe, J., Dos Santos, S. E., Dunville, K., Feldmeyer, D., Fiáth, R., Fishell, G. J., Foggetti, A., Gao, X., Ghaderi, P., Goriounova, N. A., Güntürkün, O., Hagihara, K., Hall, V. J., Helmstaedter, M., Herculano-Houzel, S., Hilscher, M. M., Hirase, H., Hjerling-Leffler, J., Hodge, R., Huang, J., Huda, R., Khodosevich, K., Kiehn, O., Koch, H., Kuebler, E. S., Kühnemund, M., Larrañaga, P., Lelieveldt, B., Louth, E. L., Lui, J. H., Mansvelder, H. D., Marin, O., Martinez-Trujillo, J., Chameh, H. M., Mohapatra, A. N., Munguba, H., Nedergaard, M., Němec, P., Ofer, N., Pfisterer, U. G., Pontes, S., Redmond, W., Rossier, J., Sanes, J. R., Scheuermann, R. H., Serrano-Saiz, E., Staiger, J. F., Somogyi, P., Tamás, G., Tolias, A. S., Tosches, M. A., García, M. T., Wozny, C., Wuttke, T. V., Liu, Y., Yuan, J., Zeng, H., & Lein, E. (2020). A community-based transcriptomics classification and nomenclature of neocortical cell types. Nature Neuroscience, 23(12), 1456–1468. https://doi.org/10.1038/s41593-020-0685-8

Zur Lage, P., Newton, F. G., & Jarman, A. P. (2019). Survey of the Ciliary Motility Machinery of Drosophila Sperm and Ciliated Mechanosensory Neurons Reveals Unexpected Cell-Type Specific Variations: A Model for Motile Ciliopathies. Frontiers in genetics, 10, 24–24. https://doi.org/10.3389/fgene.2019.00024

